# Kin Recognition in a Clonal Fish, *Poecilia Formosa*

**DOI:** 10.1101/055848

**Authors:** Amber M. Makowicz, Ralph Tiedemann, Rachel N. Steele, Ingo Schlupp

**Affiliations:** Department of Biology, Ecology and Evolutionary Biology, University of Oklahoma, 730 Van Vleet Oval, Norman, OK 73019, USA; Department of Biology, Lehrstuhl für Zoologie und Evolutionsbiologie, University Konstanz, Universitätsstraβe 10, 78457 Konstanz, Germany; Unit of Evolutionary Biology/Systematic Zoology, Institute of Biochemistry and Biology, University of Potsdam, Karl-Liebknecht-Strasse 24-25, 14476 Golm, Germany

**Keywords:** Amazon molly, clonal reproduction, female aggression, kin recognition, *Poecilia formosa*

## Abstract

Relatedness strongly influences social behaviors in a wide variety of species. For most species, the highest typical degree of relatedness is between full siblings with 50% shared genes. However, this is poorly understood in species with unusually high relatedness between individuals: clonal organisms. Although there has been some investigation into clonal invertebrates and yeast, nothing is known about kin selection in clonal vertebrates. We show that a clonal fish, the Amazon molly (*Poecilia formosa*), can distinguish between different clonal lineages, associating with genetically identical, sister clonals, and use multiple sensory modalities. Also, they scale their aggressive behaviors according to the relatedness to other females: they are more aggressive to non-related clones. Our results demonstrate that even in species with very small genetic differences between individuals, kin recognition can be adaptive. Their discriminatory abilities and regulation of costly behaviors provides a powerful example of natural selection in species with limited genetic diversity.

---

Kin selection theory predicts that cooperative and altruistic behaviors scale with relatedness [1–5], strongly favoring close relatives, and have been shown empirically in numerous sexual species [Insects 6–7; Frogs 3–4; Fish 8–10; Birds 11; Mammals 5] But how large must the difference in relatedness be for kin recognition to occur [3]? To address this, we need to understand just how relatedness shapes social behavior in species with the highest possible relatedness between individuals: clonal organisms. Like monozygotic twins in humans, clonal organisms are genetically extremely similar, sometimes completely identical. Young human twins are almost impossible to tell apart by the naïve observer, but with some experience there are often subtle differences that allow us to distinguish between individuals [12]. While some clonal invertebrates are capable of detecting and favoring full clonal sisters, others lack the ability to discriminate between their own and other clonal lineages [13–18]. Indeed, it would seem that a major form of evolutionary selection, kin selection, is eliminated because the genetic variation that allows for discrimination is so minute that discrimination becomes unlikely, or the cost of altruistic behaviors become too high (i.e. limited dispersal, increased competition among relatives) [19]. These findings raise three important questions: how much genetic variation is required for kin recognition to evolve, is selection eliminated because there are no available recognition mechanisms in clonal organisms, and what is the adaptive benefit of kin recognition among clones? We investigated these questions using an ameiotic, clonal fish, the Amazon molly (*Poecilia formosa*), which naturally occurs in mixed groups of different clones [20–21].

The Amazon molly is a natural hybrid species that reproduces via sperm-dependent parthenogenesis, or gynogenesis, that originated from a hybridization event between *P. latipinna* and *P. mexicana* approx. 120,000 generations ago and has evolved into a species with extremely limited within-species variability [20,22]. The diploid eggs of *P. formosa* are pseudo-fertilized by either male *P. latipinna* in south Texas or *P. mexicana* in east Mexico, and typically the male genome is not incorporated into the offspring, leading to identical daughter clones [21,23]. Mollies are livebearing and have internal fertilization, and sexual and asexual females compete for the same males [24]. Gynogenesis results in populations of Amazon mollies that are genetically relatively uniform, yet clonal lineages may occasionally diversify by introgression, mutation, or gene conversion [21], and several different clonal lineages are known to coexist within the same population [20–21].

Amazon mollies show great similarities with their sexual hosts in their ecological niche, including feeding behavior, mating preferences, parasite loads, life history traits, fecundity, and survivorship [25–29]. They live in very fluid social environments with their host species, which change constantly in the composition of sex, species, and even clone lineage [25]. This competition should favor targeted aggressive behaviors, which in turn should favor species and potentially kin recognition. In the present study, we test the ability of *P. formosa* to distinguish clonal sisters (i.e., females of the same clone born of the same mother) from non-sisters. We further establish the sensory systems used in this recognition, and provide an adaptive explanation for the evolution of kin recognition by testing if aggressive behaviors scale directly with relatedness.

Given the high genetic similarity with other clonal lineages, competition for resources, low dispersal rates of the clones, the high diversity of clonal lineages with in a population, and the social environment in which they occur, we hypothesize that Amazon mollies show the ability to detect different clonal lineages and adjust their aggressive behaviors accordingly.

To test this hypothesis, we created six clonal lineages by mating virgin Amazon mollies from populations collected from the entire geographical range of the species to sailfin molly males (Fig. S1, Table S1). Clonality of each lineage was confirmed using microsatellites (Tables S2, S3, S4). The results indicate that our clonal lineages exhibit: 1) high degrees of relatedness within each clonal lineage of or close to the value of 1; and 2) lower relatedness between clonal lineages in comparison (Table S5, S6). We define clonal sisters as those individuals that are genetically identical, based on microsatellites, to the focal females and are descendants of the same founding mother. Non-sister individuals are defined as females that originate from a different, more distant clonal lineage and are not genetically identical to the focal females (i.e., as related to the focal females as random, non-kin individuals of a sexual species). Additionally, Amazon mollies show considerable individual variation in behaviors (i.e., preferences, aggression, etc.) [24,30–31] within and among clonal lineages, suggesting that after establishing kin recognition in multiple lineages, the use of a single lineage to further explore kin recognition within this species is sufficient.

## Material and methods

### (a) Populations

A single female each from six populations across the geographic range of *Poecilia formosa* (Fig. S1) was isolated and kept with a male *P. latipinna* (Comal Spring, TX) to found the clonal lineages (Table S1). Populations were maintained in outdoor tanks (1000L) during the summer and indoor tanks in the winter and fed tropical fish flakes *ad libitum.* After several generations (4±2 generations), tissue samples were collected to confirm that the population was a single clonal lineage. We used 12 microsatellites to analyze the genetic divergence between the different populations (Table S2) [38]. We then compared loci, H_0_, H_E_ of each of the different clonal lineages. We also assessed divergence among lineages, by calculating the FST values among lineages, both locus-wise and across all loci, and by performing exact tests of differentiation using Markov Chain Monte Carlo simulations (Table S2, S3) [39]. Genotypes of females indicate that each female within the clonal lineages is indeed identical to one another. We calculated the genetic identity and relatedness coefficient within and among the clonal lineages (Supplementary Tables 5, 6) [40–41]. We found that although all Amazons are closely related, females clustered together based on clonal lineage. We also wanted to investigate kin recognition within a population, and isolated two different clones from Comal Spring, TX. These clones only differed at two microsatellite loci (GA-V18: 122–144 vs. 122–148; GT-II33: 182–182 (homozygous) vs. 178–182). Together this allows us to address the minimum genetic distance required for kin recognition to occur within a population and between populations. Note that one clonal lineage, Comal Spring 7a, was genetically indistinguishable from that of San Ignacio with the 12 microsatellites that we tested for (Table S4). All experiments were approved by the University of Oklahoma Animal Care and Use Committee (#R13–006).

### (b) Kin recognition

A standard binary choice test (Fig. S3) [42–43] allowed *P. formosa* to choose between clonal sisters and non-sisters (a clone from another population). The stimulus fish (size matched females, ±3mm) were placed in clear, perforated Plexiglas cylinders (to allow chemical, visual, and mechanical cues) at each end of the experimental tank (61×39×30cm; Note: these Plexiglas cylinders differed depending on the protocol to test specific types of cues) The focal females were then placed into the center of the tank inside a Plexiglas cylinder to permit chemical, visual, and mechanical cues to reach them and were then allowed to acclimate for 10 minutes. After this period, the association time (s) females spent in the preference zone with a stimulus female was recorded. The experiment did not begin until the female began swimming freely. To prevent any side bias, focal females were tested twice, with the second trial having the partner females switching sides (with exception to the mechanisms experiment, see below) [43]. These two trials were added together and the strength of preference (SOP) scores were calculated as: The total time spent with Stimulus 1 / (Total time spent Stimulus 1 + Total time spent Stimulus 2). These SOP scores were calculated for both the clonal sisters and non-sisters, then √arc(sin) transformed to normalize the data. Paired *t*-tests were used to compare the transformed SOP (Strength of Preference) scores for clonal sisters and non-sisters in SPSS (ver. 17, Fig. 1, 2).

**Figure 1.**
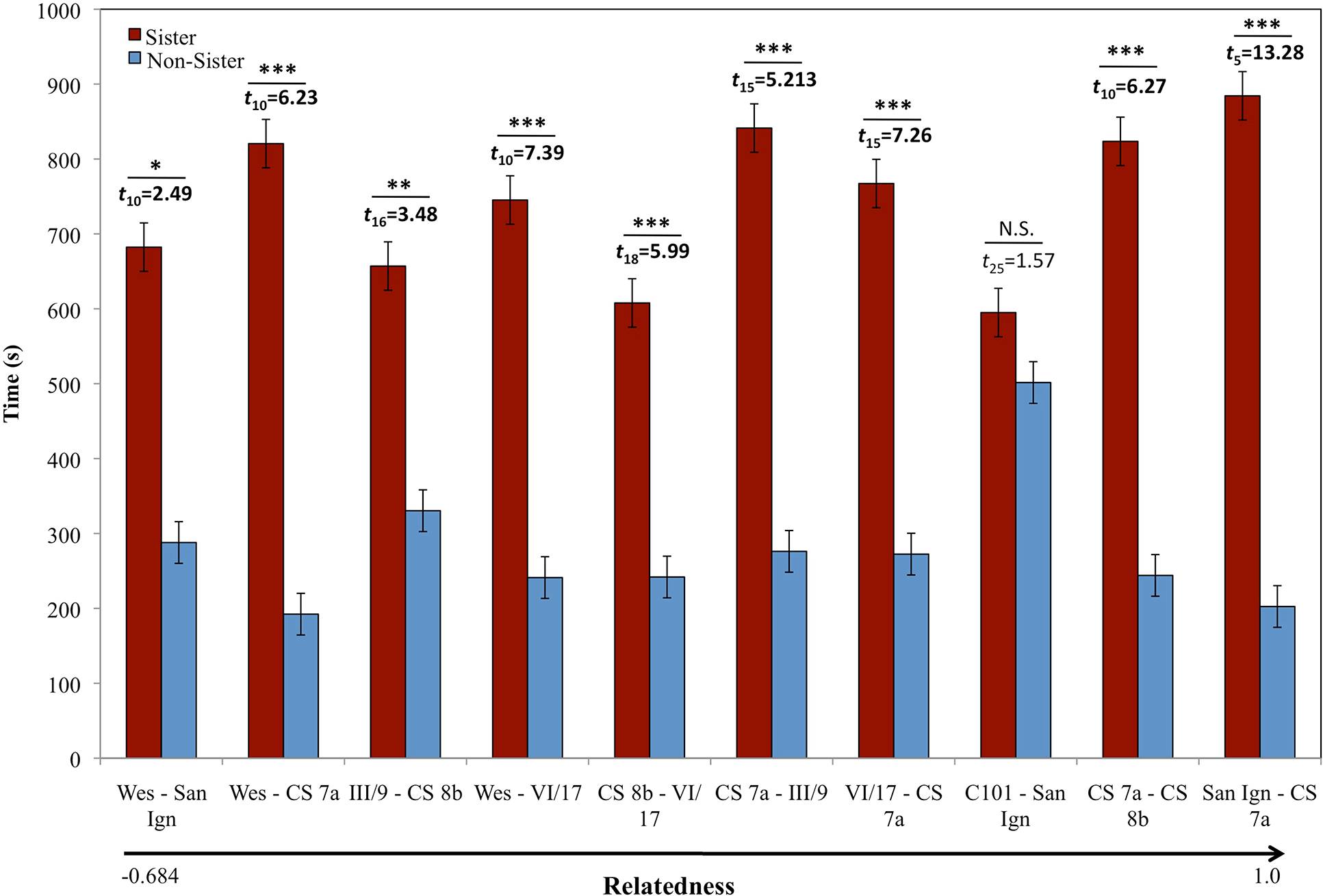
**Female Kin Preference.** The average time ± SE female preferences for clonal sisters (red) and non-sisters (blue) in six different clonal lineages across the range of P. formosa. Relatedness between the focal females and the nonsister clones scaled from left (more distant) to right (identical). Weslaco (Wes) paired with non-sister San Ignacio (San Ign); Weslaco paired with non-sister Comal Spring 7a (CS7a); III/9 Barretal (III/9) paired with non-sister Comal Spring 8b (CS8b); Weslaco paired with non-sister VI/17 Nuevo Padilla, (VI/17); Comal Spring 8b paired with non-sister VI/17 Nuevo Padilla; Comal Spring 7a paired with non-sister III/9 Barretal; VI/17 Nuevo Padilla paired with non-sister Comal Spring 7a; County 101 San Marcos (C101) paired with non-sister San Ignacio; Comal Spring 7a paired with non-sister Comal Spring 8b; and San Ignacio paired with non-sister Comal Spring 7a. Females from 6 of the 7 populations showed a significant preference (* = p<0.05; ** = p<0.009; *** = p<0.0001; NS = non-significant) for clonal sisters over non-sisters when visual, chemical and mechanical information was present. For unknown reasons, C101 clonal lineage had relatively low genetic identity, likely leading to a lack of kin recognition in this population.

**Figure 2.**
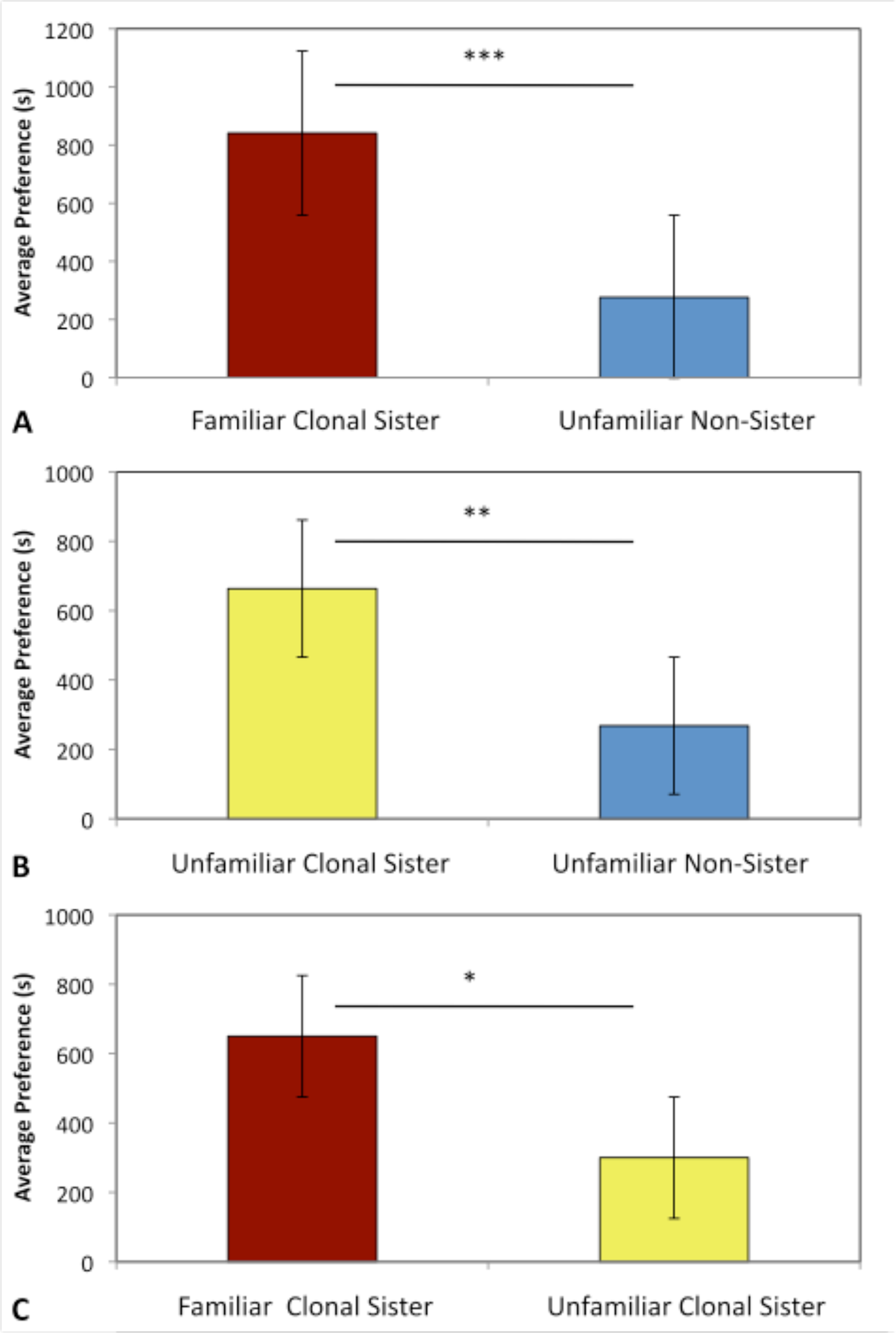
**Preferences for Familiarity.** The average ± SE for female preferences of clonal sisters and non-sisters was not due to familiarity. In each experiment, 15 females were given a choice between a familiar clonal sister (red), an unfamiliar clonal sister (yellow), or an unfamiliar non-sister (blue) after a 10-month isolation period from clonal sisters. **A.** Females from Comal Spring, TX, familiar clonal sisters and unfamiliar non-sisters, *t*_15_=5.213, *p*<**0.0001**; **B.** Unfamiliar clonal sisters and unfamiliar non-sisters, *t*_15_=3.362, *p*=**0.005**; **C.** Familiar clonal sisters and unfamiliar clonal sisters, *t*_14_=2.966, *p*=**0.011**. Females maintain the preference for clonal sisters (regardless of familiarity) and prefer familiar clonal sisters to unfamiliar clonal sisters.

### (c) Mechanism of kin recognition

Focal females and clonal sisters came from a stock population originally collected from Río Purificacíon in Nuevo Padilla (VI/17), Mexico, and non-sister stimulus females originated from Comal Springs, Texas. Fish were maintained in a 12:12 light:dark photoperiod, and fed fish flakes *ad libitum* daily. Focal females were randomly selected from the stock populations and then isolated from clonal sisters for a minimum of one-week prior to conducting the experiments in a separate 75.7L tank. Stimulus female populations were maintained under similar conditions in separate 37.9L tanks.

To prevent effects residual chemical signals on female preferences, prior to each individual experiment, the experimental tanks, Plexiglas cylinders and Plexiglas sideboards were washed with soapy water, and then rinsed thoroughly, followed by 3% hydrogen peroxide [44]. After tanks were clean, the experimental tank was filled with ionized water (700–1000ppm). White Plexiglas was placed on the bottom and both long sides of the tank to prevent any influence by the presence of the experimenter. Treatment Plexiglas cylinders were randomly assigned and placed in each end on the tank.

Experimental treatments consisted of: 1) allowing visual only cues of the stimulus females to be passed to the focal fish using solid, clear Plexiglas cylinders; 2) visual and chemical cues using clear cylinders perforated with small 3mm holes; 3) chemical only cues using solid black cylinders perforated with small 3mm holes (corresponding to an area of about 7mm^2^ per hole, to reduce neuromast stimulation); 4) only chemical and mechanical (lateral line) signals using black cylinders with large 5mm holes (equivalent to about 20mm^2^ per hole, to increase neuromast stimulation); 5) a side bias control using clear empty cylinders; and 6) a color control to compare the response effects of one empty black and a clear cylinder. The area perforated was the same for cylinders with small or large holes (see Fig. S4 for visual of experimental cylinders).

A standard binary choice test was used as described above. Individual trials were run for 10-minutes using the Viewer video tracking system (BIOBSERVE GmbH, Bonn, Germany) [45] to record the time spent near each stimulus and the number of times the focal fish entered each preference zone. At the conclusion of the trial, each female was placed into an individual tank (3.8L) overnight, which allowed us to identify each focal and stimulus female. Measurements of the water chemistry (pH, salinity, temperature, and dissolved oxygen) were taken using a Horiba water quality-measuring device (U-50 series) to confirm that the water quality remained constant (data not shown). On the next day, the same procedures were employed until all six treatments were completed. The order in which the treatments were presented and the side each stimulus female was placed was randomized. During the entire duration of the experiments females were maintained in similar lighting conditions as above, and fed frozen mosquito larvae.

SOP scores were calculated using the time spent within the preference zone, and then √arc(sin) transformed to normalize the data. A repeated-measures GLM was run using “treatments” as the within-subject variable. To analyze differences between unimodal (e.g. visual only) and bimodal (e.g. visual plus chemical) sensory mechanisms a repeated-measures GLM was used using “mode” as the within subject factor.

Prior to experimentation, we validated the construction of the Plexiglas cylinders so that they were only allowed the specific signals for each type of cylinder to be released. To demonstrate the diffusion of the chemical cues from the inside of the Plexiglas cylinders we ran several trials using food coloring. Using an identical set up as previously mentioned, we placed a fish inside the cylinder and then added 10 drops of red food coloring (Ingredients: water, propylene glycol, FD&C red, and propylparaben; Note: this was not harmful to the fish) and measured the time it took to diffuse to the preference zone. We ran this trial for all four types of Plexiglas cylinders to confirm their construction (i.e., to confirm that the solid cylinder was indeed only allowing the visual only signals and did not leak chemical signals; Fig. S4).

### (d) Diet influence on chemical recognition

Fish originated from two of the single clonal lineages above (San Ignacio and Weslaco). Pregnant females were collected from stock tanks and isolated in individual tanks (3.8L) until they had offspring. Adult females were then returned to the stock tanks. Broods were raised together for a total of five weeks in 12/12 hour light/dark cycle and fed *ad libitum* brine shrimp and flake food. At 5 weeks of age, juveniles were raised individually in 3.8L tanks on either: 1) a high protein (crude protein 52% min.) diet, 2) a low protein (crude protein 37% min.) diet, or 3) a 50/50 mix of the high protein and low protein (crude protein 44.5% min.) diet. Visual communication between the tanks was prevented to avoid visual imprinting from the neighboring tanks as the juveniles grew. Each tank had weekly 2/3 water changes. The tank temperature was maintained at 27.8°C during the duration of the experiment. Individuals were raised until 22–34 weeks old prior to the start of the behavioral experiments.

A standard binary choice test was used. However, stimulus females were placed into black perforated Plexiglas rectangular cylinders on either end of the experimental tank (18.9L). These black cylinders allowed focal females to make their choice solely based on chemical cues. After the experiment was finished the focal and stimulus females were returned to their appropriate individual tanks. We used five, randomized treatments to assess whether females would retain clonal recognition when females were placed on different diets: 1) a clonal sister on a different diet vs. non-sister on same diet; 2) a clonal sister on different diet vs. non-sister on mix diet; 3) a clonal sister on mix diet vs. non-sister on same diet; 4) a clonal non-sister on same diet vs. non-sister on different diet; and 5) a clonal non-sister on same diet vs. non-sister on mix diet. Focal females were retested every 24-hours until they complete all five treatments. If a female did not respond within the first 5 minutes of the trial, the trial was terminated, and the female was returned into her appropriate tank and retested the next day. Females that were used as stimulus females were not tested as focal females until one week had passed. Females that were focal females were used as stimulus females only after all five treatments were complete.

Shoaling preference was analyzed using a preference function test (Fig. S5). Using a block design, we randomly tested half of the females as focal females and used the other half to compose the stimulus shoals; after one week, the females were switched and the second half of the females were tested as focal female with the first half as was used as stimulus females. We used five randomized treatments, and one treatment was tested every 24 hours: 1) a clonal sister shoal on the same diet, 2) a clonal sister shoal on a different diet, 3) a non-sister clonal shoal on the same diet, 4) a non-sister clonal shoal on a different diet, and 5) a control where the black Plexiglas cylinder was present in the test tank but empty. Focal females were placed in a perforated Plexiglas cylinder on the side of the tank opposite the shoal Plexiglas and allowed to acclimate for five minutes. Once the focal female's cylinder was removed she was allowed to swim freely for ten minutes. We recorded the time (s) females spent in both the preference zone (17.8cm) and the interaction zone (included the stimulus Plexiglas cylinder plus one body length from the stimulus females, 10.5cm).

For the preference test, we calculated the SOP scores for time spent with the stimulus females. These scores were then √arc (sin) transformed to normalize the data. We used a repeated-measures GLM to compare preference scores across the different treatments, with “treatment” and “stimulus type” being the within-subject factors. We used the age of the fish at the time of testing, the population the females originated from, and whether they had the same mother (maternal effects) as covariates. These factors were non-significant and were therefore removed from the model (Age, *F*_1,20_=0.415, *p*=0.923; Population, *F*_1,18_=0.088, *p*=0.916; Mother, *F*_1,20_=1.3 3 6, *p*= 0.291). For the shoal preference function test, we also used a repeated-measures GLM with “treatment” and “zone” as within-subject factors, and with “clone type” and “diet” as between-subject factors. We used “block” as a covariate, however, this did not have a significant effect on either the type of stimulus (*F*_4,16_=1.109, *p*=0.387) or the zone (*F*_1,19_=0.215, *p*=0.648) and we removed it from the model.

### (e) Kin recognition as a means to regulate aggression

Using the same females as the above mentioned experiment investigating mechanisms, focal females were tested for their aggressive behaviors towards clonal sisters and non-sisters using two experimental designs: 1) a forced-choice (i.e., one stimulus female at a time), and 2) free-swimming with choice (i.e., both stimulus females at the same time). Fish were given one week of rest in between the two experiments. Since these fish appear identical to the human eye, focal females had half of the dorsal fin clipped for identification. Both clonal sister and non-sister females underwent the same handling procedures as the focal female, although only one of them had their caudal fin was clipped, resulting in all three females visibly distinguishable from one another. All females were allowed to rest from handling for three days, prior to any trials.

*Forced-Choice Experiment.* Aggression was measured in a direct-contact (stimulus and focal female able the directly interact with one another) experimental tank (19L) with either a clonal sister or non-sister (Fig. S6). At the start of the experiments, both focal female and stimulus female were placed in separate, clear Plexiglas cylinders. After a five-minute acclimation period females were released from the cylinders and behavioral measurements (bites, tail beats, and overall time spent being aggressive) were started at the first sign of aggression and ran for 10 minutes. We measured all three behaviors, both given to the stimulus females and received from the stimulus females. At the end of the trial both females were placed back into their individual tanks. Focal females were retested 24-hours later with the other partner, either the clonal sister or non-sister that was not tested the day before, following the same procedure. A repeated-measures GLM was employed using “Clone” and “Behavior” as the within subject factors.

*Open Field Experiment.* The open field, free-swimming aggression trials took place in a 19L experimental tank with all three females together to give the focal female a choice between the two different stimuli (Fig. S7). At the start of the experiments, both focal female and stimulus females were placed in separate clear, Plexiglas cylinders. After a 5-minute acclimation period females were released from the cylinders and behavioral measurements were started at the first sign of aggression and ran for 10-minutes. At the end of the trial all females were placed back into their individual tanks. After the completion of the experiment, females were allowed to recover and regenerate their fins. A multivariate GLM was run using “clone” as the fixed factor and the “behaviors” (bites given, tail beats given, time given, bites received, tail beats received, and time received) as the dependent variables. For both experiments, if there were no aggressive interactions among the three females after 10-minutes, the trial was terminated and the focal female was retested in 24-hours.

This research was carried out in strict accordance with the recommendations in the Guide for the Care and Use of Laboratory Animals of the National Institutes of Health. The Institutional Animal Care and Use Committee of the University of Oklahoma approved this research (#R13- 006). All efforts were made to reduce and minimize any suffering that the fish may have experienced during the course of this research.

## Results/ discussion

Using standard binary choice tests, we determined if individuals from the seven clonal lineages preferred to associate with clonal sisters to non-sisters in multiple combinations. We found that six clonal lineages exhibited a significant preference for their clonal sisters both within population lineages (CS7a - CS8b) and between population lineages (Fig. 1, Table S7), indicating that they distinguish between clonal lineages. In addition, we found no evidence to support the phenotypic matching hypothesis (i.e., the strength of discrimination does not correlate with the genetic similarity between clonal lineages, Fig. 1). To determine whether this result was due to familiarity, we split sisters from one clone (CS7a) in two groups under the same conditions for over nine months (average life expectancy is 1–3 years, and sexual maturity is reached around 3 months of age) and tested the offspring of these individuals, also using standard choice tests, for their ability to recognize clonal sisters. If the recognition mechanism was based on familiarity, females should be unable to recognize unfamiliar clonal sisters. We found that females preferred clonal sisters that were unfamiliar to non-sisters (*t*_15_=3.362, *p*=0.005), and familiar clonal sisters to unfamiliar clonal sisters (*t*_14_=2.966, *p*=0.011; Fig. 2). This result indicates that familiarity is not necessary for clonal recognition, but may strengthen the preference. Therefore, we hypothesize that a genetically based recognition mechanism for phenotype matching is adaptive for Amazon mollies. We were able to confirm our findings in a field experiment, in which wild Amazon mollies from their site of origin (Weslaco), in natural water, were allowed to choose between wild caught individuals and non-sisters from VI/17 (R = −0.264), a distant laboratory lineage. We found that wild caught females retain the ability to discriminate between clonal sisters and non-sisters in natural water (data not shown).

Amazon mollies show clear preferences for clonal sisters, but which sensory information is used to assess clonal identity? We concentrated on visual, chemical, and tactile information, all of which has been shown to be important in livebearing fishes. Using a repeated-measures design, we tested what cue or combination of cues (visual only, chemical only, visual and chemical cues, and chemical and mechanical cues) might be used by Amazon mollies to distinguish clonal sisters from non-sisters. All sensory modalities in isolation and in combination were sufficient for kin recognition, although there was no significant difference among sensory modalities (Mechanism: *F*_3,15_=0.955, *p*=0.439; Fig. 3). Within each modality, post-hoc analyses indicate that females showed the strongest preference for clonal sisters when only visual cues were present; nonetheless, they still showed a significant preference when only chemical cues, a combination of chemical/mechanical, or visual/chemical cues were presented. Female activity, however, was higher when chemical cues were present, and they entered the preference zones that included the clonal sisters more often (F_1,17_=8.285, *p*=0.010). Although it is known that Amazons prefer conspecific females when compared to their heterospecific host even when chemical only cues are present [30], here we show that their discriminatory abilities are even more precise than previously thought. In addition, we found no difference in the strength of kin recognition in the presence of unimodal and bimodal cues (F_1,70_=1.256, *p*=0.266), suggesting that discrimination is not improved using more than one sensory channel. This lends support to the conclusion that signals are often redundant, conveying comparable cues [32]. Most importantly, we were able to find the same effect in natural water. As in the laboratory, wild caught females preferred clonal sisters when chemical information was available, while the preference using visual information was not detectable (data not shown). This was likely due to naturally high turbidity of the water [33] as Amazon mollies are found in both turbid and clear environments and it is possible that they may rely more on either visual or chemical cues depending on the environment they live in.

**Figure 3.**
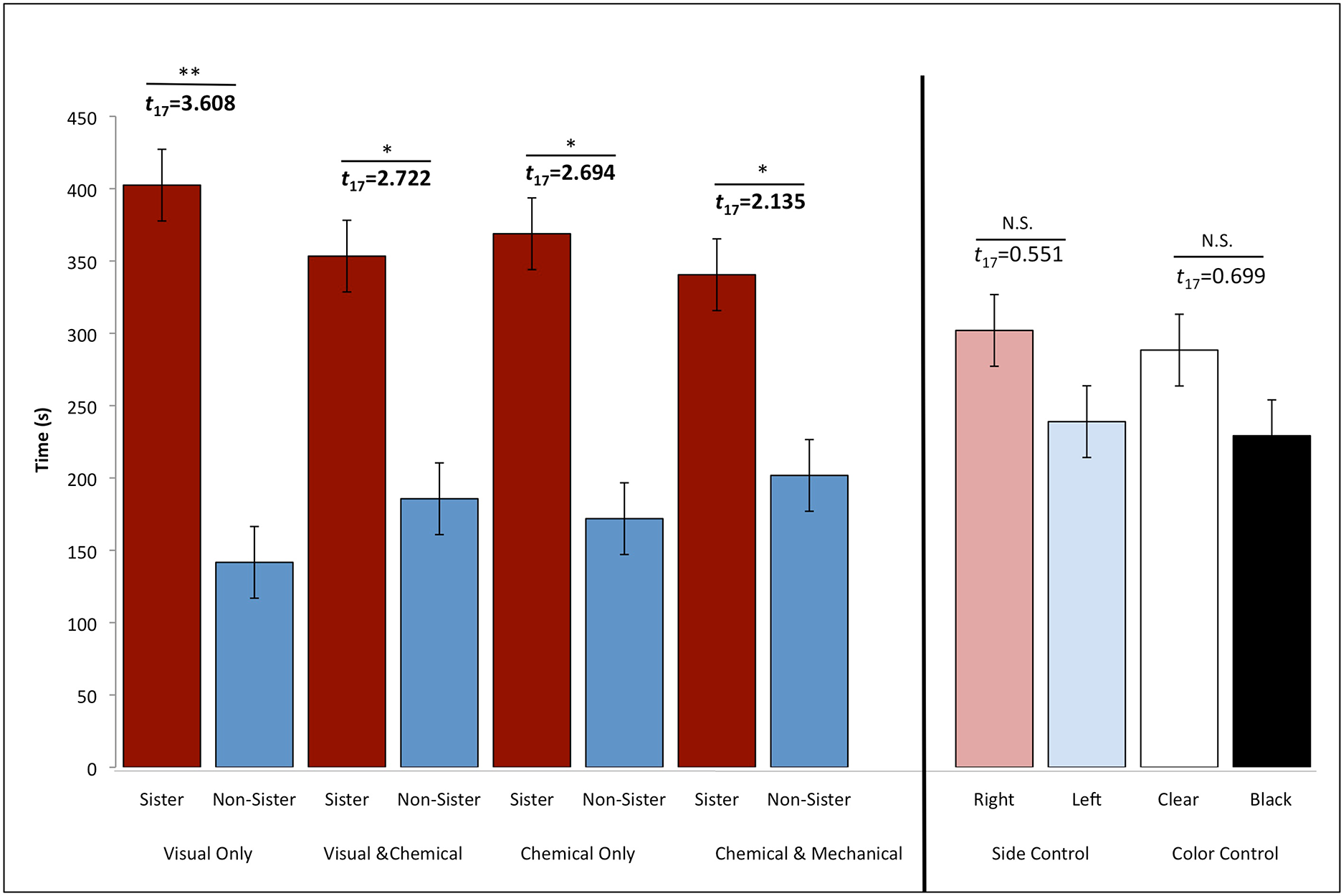
**Mechanism Experiment.** The average time ± SE females spent with a clonal sister (red; VI/17) and a non-sister (blue; CS7a; R = −0.057) in the four different treatments. Females showed a stronger preference to only visual signals (* = p<0.05; ** = p<0.009; NS = nonsignificant). The two control treatments demonstrate that there was no bias for the right (light red) or left (light blue) sides of the experimental tank and there was no bias for the clear (white) cylinder or the black (black) cylinder.

Nonetheless, there are various visual (i.e., body shape, pigment cell quantity and expression, etc.) and chemical cues (dietary, MHC genes, maternally inherited micro-biomes, etc.) in which clones may differ. Using geomorphometric analysis, we investigated body shape as a potential visual cue, and found females from clone CS7a to be significantly different from the females from clone VI/17 in body shape (data not shown). Overall, Amazon females from clone CS7a had deeper bodies, a more terminal mouth, a larger head, and a slightly longer and deeper caudal-peduncle. Body symmetry, however, did not differ between the clonal lineages. For a potential chemical signal, we evaluated how diets may influence individual preference via chemical only cues using a common garden experimental design. We found that inexperienced females retain a preference for clonal sisters on a different diet over non-sisters on the same diet (*t*_33_=3.643, *p*=0.001) and prefer to spend more time interacting with clonal sister shoals, regardless of the diet they were on, as compared to non-sister shoals (Fig. 4). This suggests that diet alone is not sufficient to alter kin recognition in these fish.

**Figure 4.**
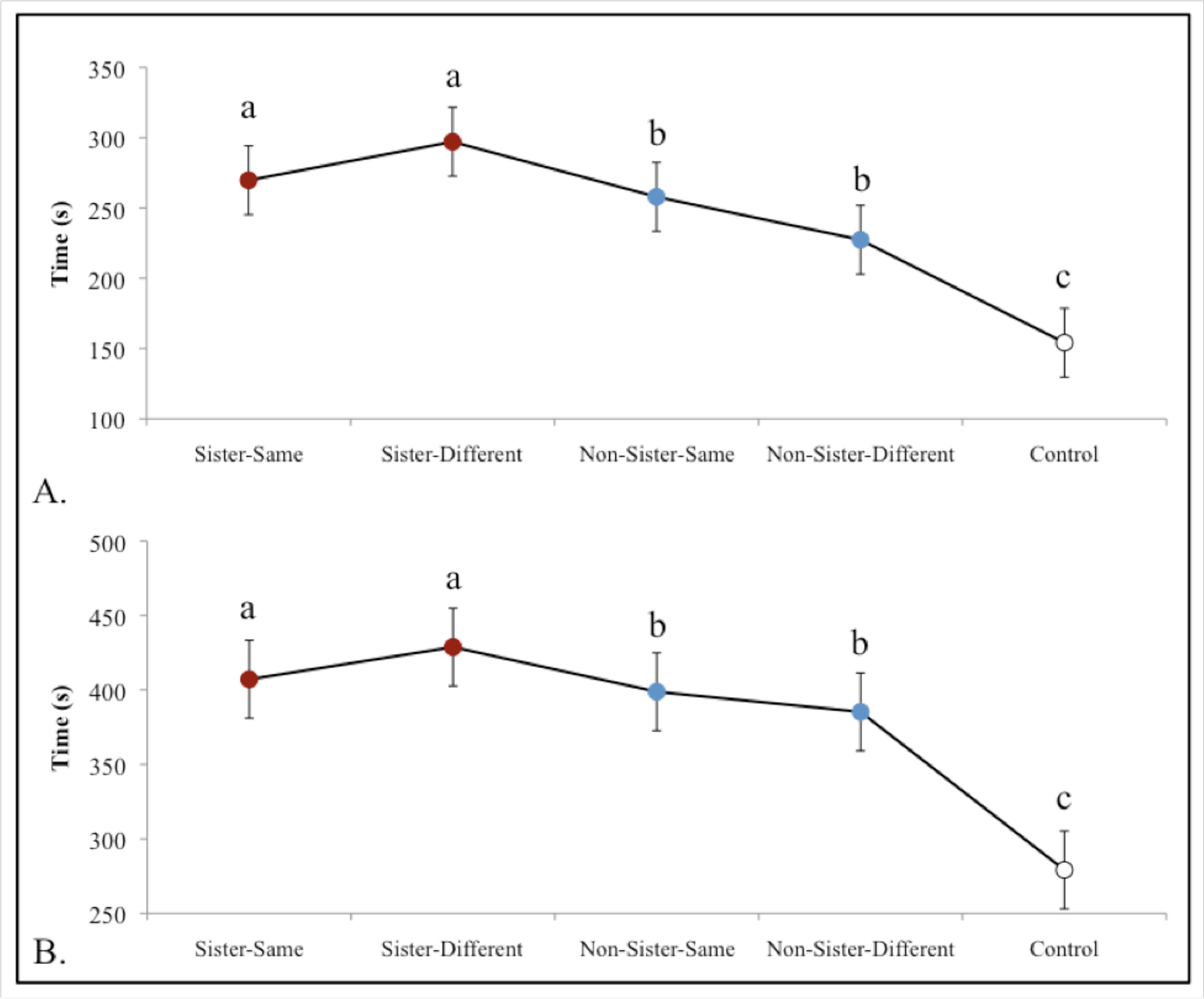
**Shoaling Preference Experiment.** Average time (s) ± SE females spent near the stimulus females in (A.) the interacting zone and (B.) the preference zone. Females spent significantly more time with the clonal sisters (red) on a different diet in both zones (preference zone: t23=2.792, p=0.010; interaction zone: t23=2.909, p=0.008) when compared to non-sisters (blue) on a different diet. They also tended to spend more time with clonal sisters on a different diet to non-sisters on the same diet (preference zone: t23=2.027, p=0.054), and with clonal sisters in general, regardless of diet, that with non-sisters (interaction zone: t47=2.132, p=0.038).

The presence of clonal recognition in an asexual vertebrate is interesting in itself, but a key question is: what adaptive benefit might Amazon mollies derive from kin recognition? Due to intraspecific competition and the extensive niche overlap between Amazons and their sexual hosts, we hypothesized that females may show more aggression towards non-sisters (and heterospecific sexual females) than clonal sisters to acquire access to limited resources, like food and potential mates [24,34–35]. Indeed, aggression in Amazon mollies has been shown to decrease their overall fitness via lower body fat condition and increasing energy expenditure [31]. We designed an open field experiment measuring the aggressive behaviors of females that were allowed to interact with both a clonal sister and non-sister. Females behaved more aggressively towards non-sisters (*F*_6,29_=2.490, *p*=0.046; Fig. 5), as would be predicted if clonal recognition is used in regulation of aggression. We also conducted a forced-choice experiment where females were allowed to interact with either a clonal sister or non-sister, which showed similar results (*F*_1,17_=8.981, *p*=0.002; Fig. S2). Together, this suggests that it is adaptive for Amazon mollies to regulate their aggressive behaviors towards clonal sisters and non-sisters due to the high cost incurred on their fitness via reduced body conditioning and potential energy available to invest into future offspring [31].

**Figure 5.**
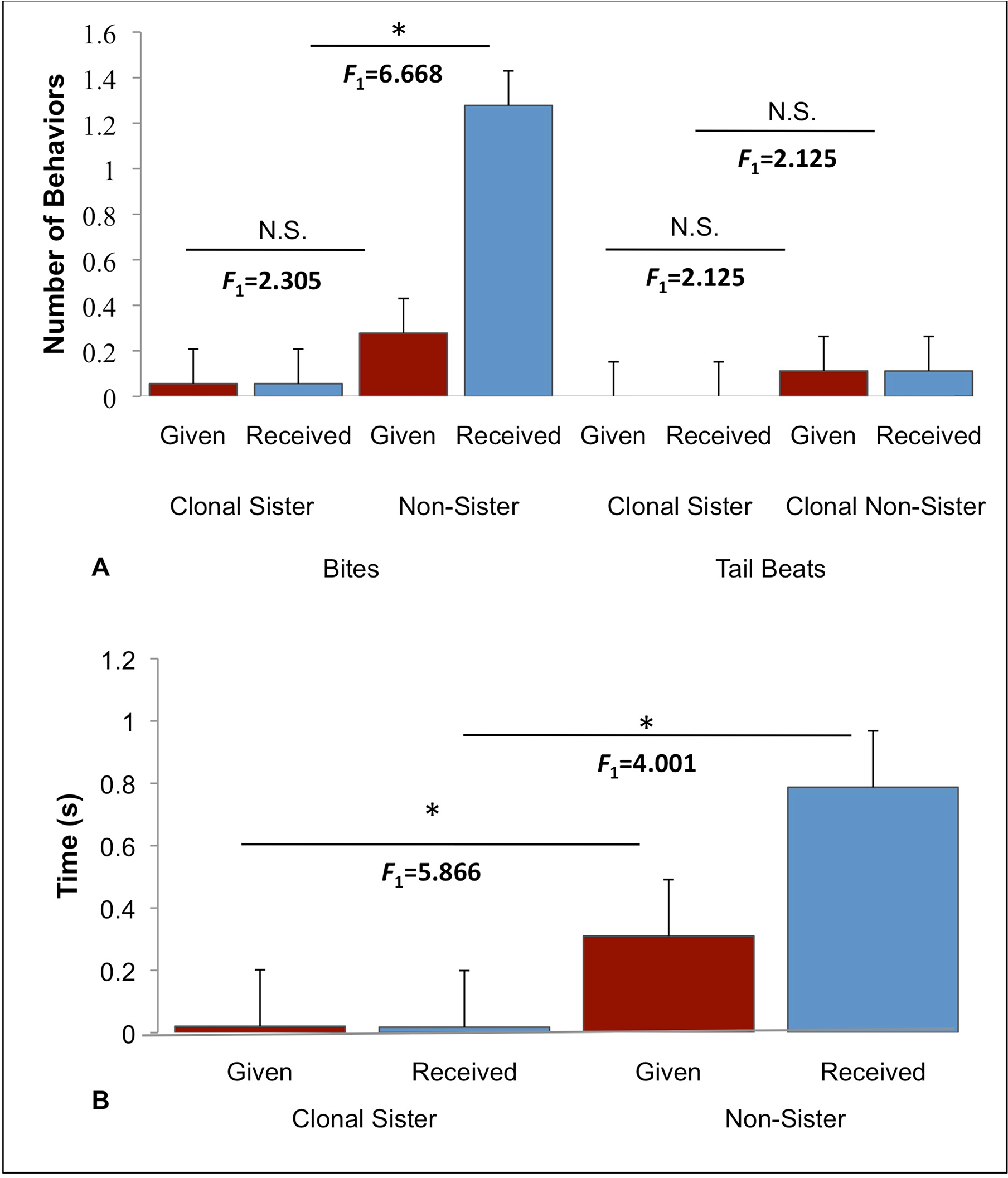
**Aggression Experiment.** This experiment tested the aggression levels of females when given a choice between a clonal sister (VI/17 Rio Purification, Nuevo Padilla, MX) and non-sister (Comal Spring, TX (7a), (Relatedness coefficient= −0.057; average±SE). Females received (blue) significantly more bites (A. given: F(1)=2.305, p=0.138; received: F(1)=6.668, p=0.014) and spent significantly more time performing (given= red) aggressive behaviors (B. given: F(1)=5.866, p=0.021; received: F(1)=4.001, p=0.054) towards non-sisters when compared to clonal sisters. There was no significant difference in performing tail beats (A. given: F(1)=2.125, p=0.154; received: F(1)=2.125, p=0.154).

In sexual species, kin recognition can evolve between closely related and distantly related individuals that are more genetically distinct (i.e., individuals share either 50% or 25% of genes that are identical by descent). It is likely to evolve when siblings overlap in time and space, and are able to recognize each other independently of context and familiarity [3]. The same parameters would hold true for asexual species; however, there is a much smaller genetic difference between individuals, which may suggest relaxed selection on other preferences. Alternatively, with female clones being genetically identical to one another, females may be using self-referential phenotype matching (using one's own cues as a reference in discriminating between kin and non-kin), which would perceptually be an easier task as compared to a sexual species. Although we did find that familiarity was present but not necessary, this may suggest that there are multiple cues in which lead to more precise discrimination between clonal sisters and non-sisters. Other factors such as maternal or epigenetic effects may also contribute to the phenotypic diversification between the different clonal lineages. However, given the controlled environment in which all the females were raised and maintained, this is an unlikely explanation for our results. Although it is known that diet and gut microbiota may influence the ability to recognize kin [36–37], we raised juveniles on different diets, manipulating their chemical signals and the gut microbiota, and found that juveniles were still able to recognize and prefer clonal sisters regardless of diet type and maternal effects. Nonetheless, epigenetics are of great interest when investigating clonal species.

With their ability for clonal recognition, Amazon mollies are one of the most extreme examples corroborating the predictions of kin selection theory, where aggression is regulated in a way that extremely close (i.e., genetically identical) kin are favored over very close kin. Given the substantial genetic similarity found throughout the whole species [20–21], up to 15 genetically distinct clonal lineages have been found within a single population with each lineage varying in frequency [AM Makowicz, unpublished data]. Nonetheless, this indicates that it is likely beneficial for clones to be able to recognize each other and regulate competition in a way that favors extremely close kin; even minute genetic differences provide enough substrate for kin recognition. We believe that the discrimination ability found in Amazons could be a powerful example of natural selection in action.

## Author contributions

This study was originally conceived by AMM and IS. AMM collected the data and implemented the data analyses with assistance from RT and RNS. AMM and IS wrote the manuscript with input from RT and RNS. Financial support for this study was obtained by AMM, IS, and RT. All authors read and approved the manuscript.

## Acknowledgements

We thank E Marsh-Matthews, G Wellborn, Z Trachtenberg, J Cureton II, M Sapage, M Tobler, D McLennan, MJ Ryan, and MA Eodice for comments on previous versions; D McLennan for methodology input; A Schneider for microsatellite genotyping; E Remmel, M Marcus, and J Nigh for data collection assistance; C Atkinson for GIS arc map assistance; and G Martin for constructing the Plexiglas cylinders. This research is in partial fulfillment of PhD dissertation requirements for AMM.

## Supporting information

### S1 Text document. Supplementary Tables And Figures

Table S1. Population origins.

Table S2. Characteristics of the 12 microsatellites.

Table S3. Genetic divergence.

Table S4. Loci difference and geographic range.

Table S5. Genetic Identity.

Table S6. Relatedness Coefficient.

Table S7. Kin Preference Scores From Multiple Populations and Pairings. Fig. S1. Map of Population Localities.

Fig. S2. Aggression Experiment #1: Forced Choice.

Fig. S3. Female Preference Experimental Set-up.

Fig. S4. Plexiglas Cylinder Construction and Functions.

Fig. S5. Shoaling Preference Experimental Tank.

Fig. S6. Female Aggression Experimental set-up #1: Forced Choice.

Fig. S7. Female Aggression Experimental set-up #2: Free-swimming with Choice.

## References

1. Hamilton WD. The evolution of altruistic behavior. Am Nat. 1963. 97, 354–356. doi: 10.1086/497114

2. Gardner A, Griffin AS, West SA. Altruism and cooperation. In Evolutionary Behavioral Ecology, (eds Westneat DF, Fox CW) New York City, NY: Oxford University Press. 2010. pp. 308–326.

3. Waldman B. The ecology of kin recognition. Annu Rev Ecol Syst. 1988. 19, 543–571.

4. Pfennig DW, Reeve HK, Sherman PW. Kin recognition and cannibalism in spadefoot toad tadpoles. Anim Behav. 1993. 46, 87–94. doi: 10.1006/anbe.1993.1164

5. Mateo JM. Kin recognition in ground squirrels and other rodents. J Mammal. 2003. 84, 1163–1181. doi: 10.1644/BLe-011

6. Breed MD, Stiller TM, Moor MJ. The ontogeny of kin discrimination cues in the honey bee, Apismellifera. Behav Genet. 1988. 18, 439–448. doi: 10.1007/BF01065513

7. Giron D, Dunn DW, Hardy ICW, Strand MR. Aggression by polyembryonic wasp soldiers correlates with kinship but not resource competition. Nature. 2004. 430, 676–679. doi: doi: 10.1038/nature02721

8. Russell ST, Kelley JL, Graves JA, Magurran AE. Kin structure and shoal composition dynamics in the guppy, Poecilia reticulata. OIKOS. 2004. 106, 520–526. doi: 10.1111/j.0030-1299.2004.12847.x

9. Gerlach G, Lysiak N. Kin recognition and inbreeding avoidance in zebrafish, *Danio rerio,* is based on phenotype matching. Anim Behav. 2006. 71, 1371–1377. doi: 10.1016/j.anbehav.2005.10.010

10. Piyapong C, Butlin RK, Faria JJ, Scruton KJ, Wang J, Krause J. Kin assortment in juvenile shoals in wild guppy populations. Heredity. 2011. 106, 749–756. doi: 10.1038/hdy.2010.115

11. Pravosudova EV, Grubb Jr. TC, Parker PG. The influence of kinship on nutritional condition and aggression levels in winter social groups of tufted titmice. Condor. 2001. 103, 821–828. doi: 10.1650/0010-5422(2001)103[0821:TIOKON]2.0.CO;2

12. Srinivas N, Aggarwal G, Flynn PJ, Vorder Bruegge RW. Analysis of facial marks to distinguish between identical twins. IEEE Trans Inf Forensics Security. 2012. 7, 1536–1550. doi: 10.1109/TIFS.2012.2206027

13. Strassmann JE, Queller DC. Selfish responses by clone invaders. Proc Natl Acad Sci USA. 2001. 98, 11839–11841. doi: 10.1073/pnas.221450998

14. Miller DG III., Consequences of communal gall occupation and a test for kin discrimination in the aphid Tamalia coweni (Cockerell)(Homoptera: Aphididae). Behav Ecol Sociobiol. 1998. 43, 95–103. doi: 10.1007/s002650050471

15. Winsor GL, Innes DJ. Sexual reproduction in Daphnia pulex (Crustacea: Cladocera): observations on male mating behaviour and avoidance of inbreeding. Freshwater Biol. 2002. 47, 441–450. doi: 10.1046/j.1365-2427.2002.00817.x

16. Segoli M, Keasar T, Harari AR, Bouskila A. Limited kin discrimination abilities mediate tolerance toward relatives in polyembryonic parasitoid wasps. Behav Ecol. 2009. 20, 1262–1267. doi: 10.1093/beheco/arp125

17. Kronauer DJC, Tsuji K, Pierce NE, Keller L. Non-nest mate discrimination and clonal colony structure in the parthogenetic any Cerapachyus biroi. Behav Ecol. 2013. 24, 617–622. doi: 10.1093/beheco/ars227

18. Loughry WJ, McDonough CM. The nine-banded armadillo. Norman, OK: University of Oklahoma Press. 2013.

19. Cornwallis CK, West SA, Griffin AS. Routes to indirect fitness in cooperatively breeding vertebrates: kin discrimination and limided dispersal. J Evolution Biol. 2009. 22, 2445–2457.

20. Stock M, Lampert KP, Möller D, Schlupp I, Schartl M. Monophyletic origin of multiple clonal lineages in an asexual fish (Poecilia formosa). Mol Ecol. 2010. 19, 5204–5215. doi:10.1111/j.1365-294X.2010.04869.x

21. Schlupp I, Riesch R. Evolution of unisexual reproduction. In Ecology and Evolution of Poeciliid Fishes (eds Evans J, Pilastro A, Schlupp I) Chicago, IL: University of Chicago Press. 2011. pp. 50–58.

22. Schartl M, Wilde B, Schlupp I, Parzefall J. Evolutionary orgin of a parthenoform, the Amazon. molly, Poecilia formosa, on the basis of a molecular genealogy. Evolution. 1995. 49, 827–835. doi: 10.2307/2410406

23. Hubbs CL, Hubbs LC. Apparent parthenogenesis in nature in a form of fish of hybrid origin. Science. 1932. 76, 628–630. doi: 10.1126/science.76.1983.628

24. Marler CA, Ryan MJ. Origin and maintenance of a female mating preference. Evolution. 1997. 51, 1244–1248. doi: 10.2307/2411053

25. Schlupp I. Behavior of fishes in the sexual/unisexual mating system of the Amazon molly (Poecilia formosa). Adv Stud Behav. 2009. 39, 153–183. doi: 10.1016/S0065-3454(09)39005-1

26. Fischer C, Schlupp I. Feeding rates in the sailfin molly Poecilia latipinna and its coexisting sexual parasite, the gynogenetic Amazon molly Poecilia formosa. J Fish Biol. 2010. 77, 285–291. doi: 10.1111/j.1095-8649.2010.02672.x

27. Tobler M, Wahli T, Schlupp I. Comparison of parasite communities in native and introduced populations of sexual and asexual mollies of the genus Poecilia. J Fish Biol. 2005. 67, 10721082. doi: 10,1111/j.1095-8649.2005.00810.x

28. Riesch R, Plath M, Makowicz AM, Schlupp I. Behavioural and life-history regulation in a unisexual/ bisexual mating system: does male mate choice affect female reproductive life histories? Biol J Linn Soc. 2012. 106, 598–606. doi: 10.1111/j.1095-8312.2012.01886.x

29. Tobler M, Schlupp I. Differential susceptibility to food stress in neonates of sexual and asexual mollies (Poecilia, Poeciliidae). Evol Ecol. 2010. 24, 39–47. doi: 10.1007/s10682-008-9288-7

30. Reding L, Cummings ME. Does sensory expansion benefit asexual species? An olfactory discrimination test in Amazon mollies. Behav Ecol. 2016. doi:10.1093/beheco/arv168.

31. Partan SR, Marler P. Issues in the classification of multimodal communication signals. Am Nat. 2005. 166, 231–245. doi: 10.1086/431246

32. Heubel KU, Schlupp I. Turbidity affects association behaviour in male Poecilia latipinna. J Fish Biol. 2006. 68, 555–568. doi: 10.1111/j.0022-1112.2006.00941.x

33. Rosvall K. Intrasexual competition in females: evidence for sexual selection? Behav Ecol. 2011. 22, 1131–1140. doi: 10.1093/beheco/arr106

34. Stockley P, Bro-J0rgensen J. Female competition and its evolutionary consequences in mammals. Biol Rev. 2011. 86, 341–366. doi: 10.1111/j.1469-185X.2010.00149.x

35. Makowicz AM, Schlupp I. Effects of female-female aggression in a unisexual/sexual species complex. Ethology. 2015. 121, 904–914. doi: 10.1111/eth.12406

36. Lize A, McKay R, Lewis Z. Kin recognition in Drosophila: the importance of ecology and gut microbiota. ISME J. 2014. 8, 469–477. doi: 10.1038/ismej.2013.157

37. Rajakaruna RS, Brown JA. Effects of dietary cues on kin discrimination in juvenile Atlantic salmon (Salmo salar) and brook trout (Salvelinus fontinalis). Can J Zool. 2006. 84, 839–845. doi:10.1139/Z06-069

38. Tiedemann R, Moll K, Paulus B, Schlupp I. New Microsatellite loci confirm hybrid origin, parthenogenetic inheritance, and mitotic gene conversion in the gynogenetic Amazon molly (Poecilia formosa). Mol Ecol. 2005. 5, 586–589. doi: 10.1111/j.1471-8286.2005.00993.x

39. Plath M, Hauswaldt JS, Moll K, Tobler M, De Leon FJG, Schlupp I, Tiedemann R. Local adaptation and pronounced genetic differentiation in an extremophile fish, Poecilia mexixcana, inhabiting a Mexican cave with toxic hydrogen sulphide. Mol Ecol. 2007. 16, 967–976. doi: 10.1111/j.1365-294X.2006.03212.x

40. Lynch M, Ritland K. Estimation of pairwise relatedness with molecular markers. Genetics. 1999. 152, 1753–1766.

41. Tiedemann R, Paulus KB, Havenstein K, Thorstensen S, Petersen A, Lyngs P, Milinkovitch MC. Alien eggs in duck nests: brood parasitism or a help from Grandma? Mol Ecol. 2011. 20, 3237–3250. doi: 10.1111/j.1365-294X.2011.05158.x

42. Schlupp I, Marler C, Ryan MJ. Benefit to male sailfin mollies of mating with heterospecific females. Science. 1994. 263, 373–374. doi: 10.1126/science.8278809

43. McCoy E, Syska N, Plath M, Schlupp I, Riesch R. Mustached males in a tropical poeciliid fish: emerging female preference selects for a novel male trait. Behav Ecol Sociobiol. 2011. 65, 1437–1445. doi: 10.1007/s00265-011-1154-x

44. McLennan DA, Ryan MJ. Interspecific recognition and discrimination based upon olfactory cues in northern swordtails. Evolution. 1999. 53, 880–888. doi: 10.2307/2640728

45. Schwarz S, Hofmann MH, Guzmen C, Schlax S, von der Emde G. VIEWER: a program for visualizing, recording, and analyzing animal behavior. Comput Meth Prog Bio. 2002. 67, 55–66. doi: 10.1016/S0169-2607(00)00150-4

